# Loss or redistribution? A better way of estimating regional changes in animal distribution and numbers caused by increased human activities

**DOI:** 10.1101/2024.09.04.611199

**Authors:** Moritz Mercker, Verena Peschko, Kai Borkenhagen, Nele Markones, Henriette Schwemmer, Volker Dierschke, Stefan Garthe

**Affiliations:** Bionum GmbH, Consultants in Ecological Statistics, Hamburg, Germany; Institute of Applied Mathematics (IAM) and Interdisciplinary Center of Scientific Computing (IWR), Heidelberg University, Heidelberg, Germany; Research and Technology Centre (FTZ), University of Kiel, Büsum, Germany; Federation of German Avifaunists (DDA), Münster, Germany

## Abstract

1. A differentiated understanding of how regional human activities affect the spatial distribution and numbers of animals within specific areas of interest is of great ecological importance. Estimating these effects from empirical data is challenging however, because human activities can affect animals in qualitatively different ways and on different spatial and temporal scales. In addition, spatio-temporal animal abundance is frequently influenced by factors intrinsic and extrinsic to the area of interest, potentially confounding impact studies, e.g., based on trends.
2. In this study, we synergistically combined regression and mechanistic modelling to separate these different influences. We first used partial differential equations to simulate various potential animal redistribution patterns affected by regional human activities. We then selected appropriate patterns as predictors in regression-based species distribution models, together with additional anthropogenic and natural covariates. The simultaneous consideration of large-scale (number-conserving) animal reorganisation, their regional loss or gain, and the influence of additional environmental covariates eventually allowed the generation of qualitative and quantitative estimates and predictions of human-induced changes.
3. We exemplarily applied our approach to investigate the current and future impact of increasing offshore wind farm (OWF) implementation in the German North Sea on common murres (*Uria aalge*) during autumn. OWFs constructed up to 2019 reduced common murre numbers in German waters by 18.3%. If the planned OWF priority and reservation areas outlined in the German Marine Spatial Plan are implemented, the predicted loss would increase to 77.7%. Notably, these predictions did not include additional anthropogenic activities or further plans for OWF installation, which could together lead to the almost complete disappearance of common murres from the German North Sea.
4. By directly comparing predicted animal numbers and distributions in hypothetical scenarios with and without human pressures, the presented method allows us to measure and predict the effects of human activities on regional trends and large-scale reorganisation. This in turn helps us to quantify and predict the impact of planned human activities on wildlife, including in the context of the current rapid expansion of alternative energies.

## Introduction

Anthropogenic activities frequently lead to distinct changes in wildlife population numbers and/or spatial patterns, and the robust quantification and prediction of these changes provides an important basis for ecosystem management [1]. Human activities can affect animals qualitatively in various ways and on different spatio-temporal scales (Fig. 1 and 2A). For example, animals within an area of interest might rearrange spatially (e.g., avoiding these activities on a meso-scale, in the offshore-scenario exemplarily considered in this study referring to a scale of kilometres), while their overall numbers may remain constant [2]. However, if there is a lack of suitable and undisturbed habitats, this redistribution may displace birds on a macro-scale across the boundaries of the study area, effectively causing a loss of individuals from the considered area. Beyond these displacements, individuals may also be completely lost from the wider region, either by migrating to alternative distant regions or as a result of increased mortality. The latter may occur as a result of a direct reduction (e.g., by collision with anthropogenic structures [3]), or by secondary effects, usually over longer time scales (e.g., as a result of reduced prey availability [4] or increased stress [5], leading to reduced survival or reproduction [6]).

**Figure 1.**
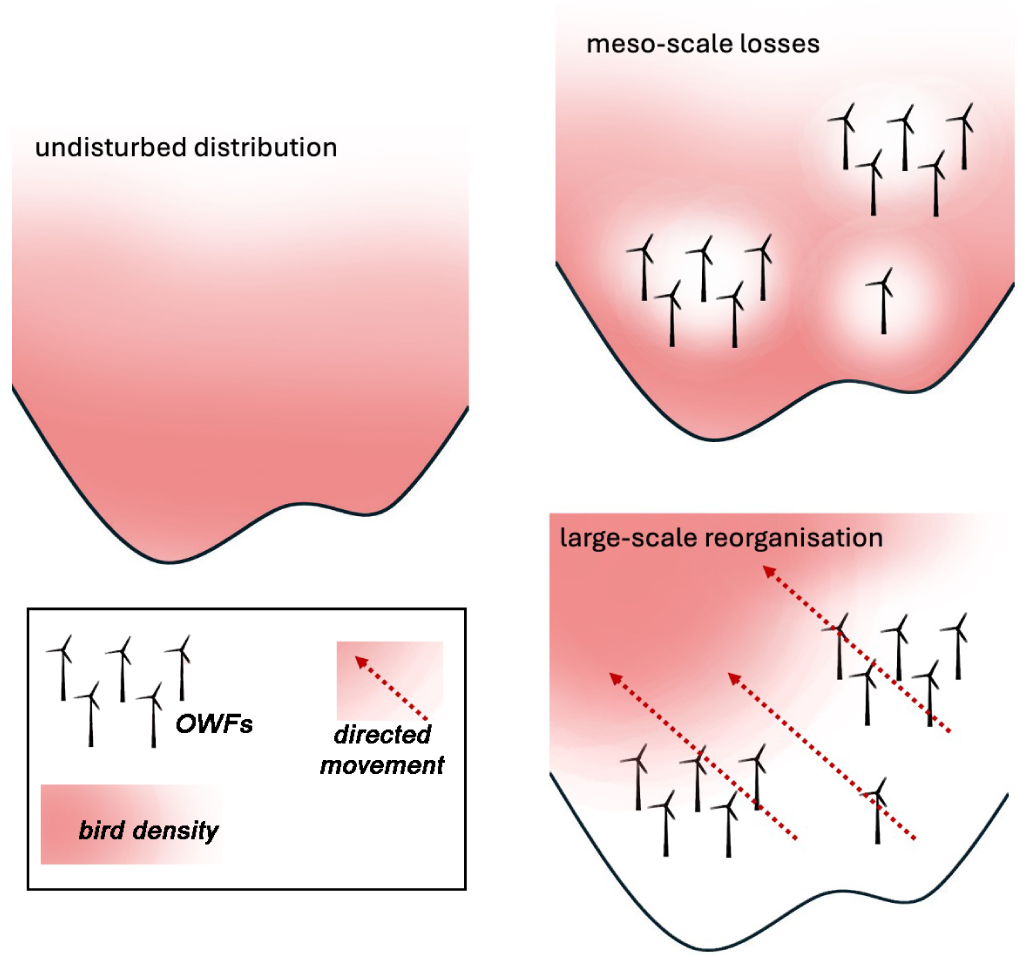
Two distinct observed reactions of seabirds on OWFs are meso-scale losses and large-scale reorganisation.

**Figure 2.**
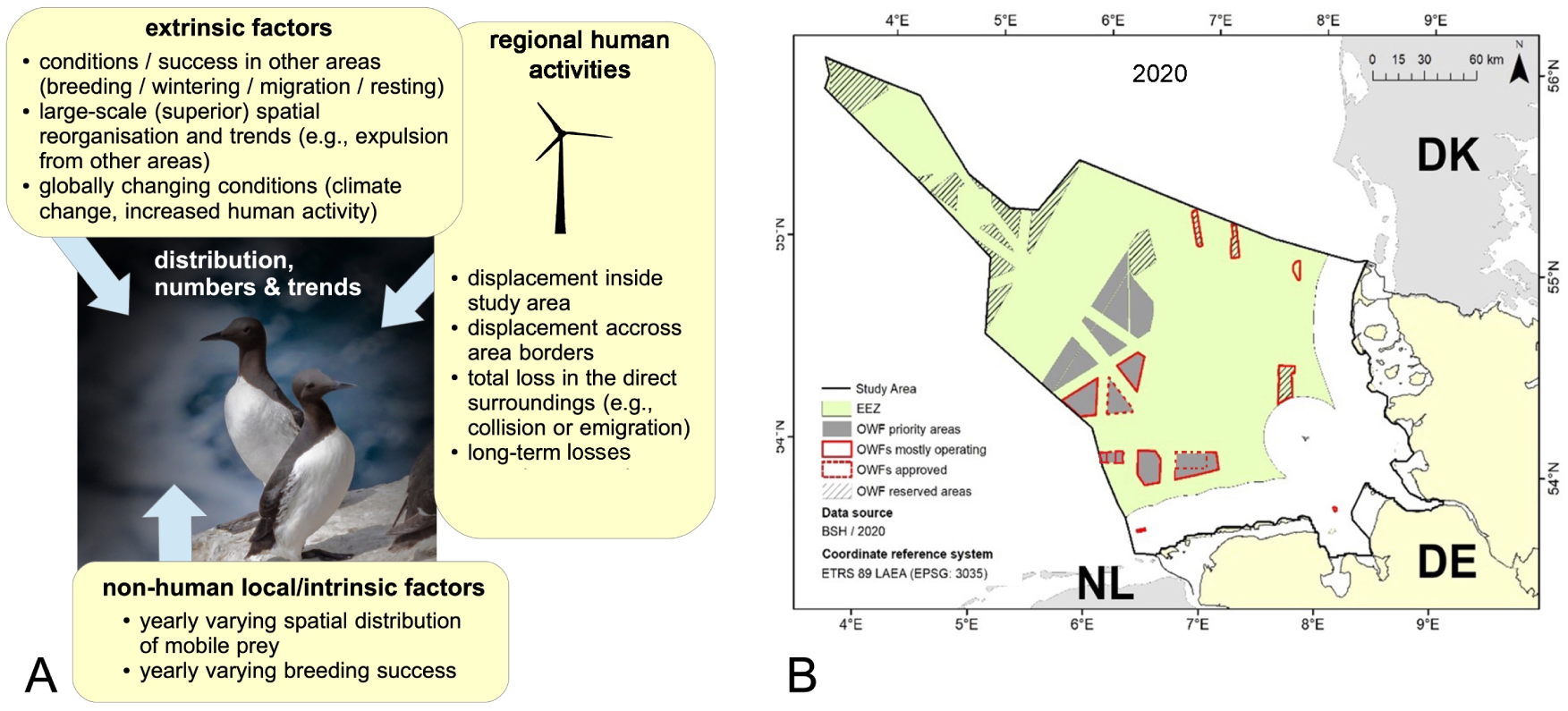
(A) Examples of factors intrinsic and extrinsic to the study area that could influence regional densities, spatial patterns, and trends in species such as common murres in the German North Sea. (B) Study area (German exclusive economic zone (EEZ) and coastal areas – black line) including operating OWFs (red-framed areas), planned (grey-shaded areas), and optional/reserved (grey-striped areas) for OWF construction. Figure modified from [28].

In addition to specific human activities, several other factors intrinsic or extrinsic to the study area may influence species distribution and numbers, thus impeding impact analyses (Fig. 2A). For example, animals can redistribute spatially within the study area depending on varying prey availability [7], leading to high spatial variability and making it difficult to interpret yearly spatial patterns. In addition, particularly for migratory species (i.e., open populations), yearly numbers within the considered area may also depend strongly on extrinsic conditions (e.g., in wintering or breeding areas), such as annual variations in reproductive success in breeding grounds, or stochastic or systematic large-scale spatial variations. Overall, especially for migratory species, trend analyses appear to be an inappropriate tool for estimating the partial effect of regional human pressures on animal numbers within an area of interest, given that trends can be largely affected by additional causes, external or instrinsic to the study area, which may strongly confound the conclusions. For example, an externally caused positive population trend (e.g., caused by spatial displacement into the study area) could be locally damped by the simultaneous intensification of human activities in the area of interest, wrongly suggesting a lack of negative effects caused by these activities.

Two main (and distinctly different) theoretical tools have been developed and applied to investigate and model the influence of human pressures on spatio-temporal animal patterns in the last decades, namely correlative/regression-based [8] and mechanistic [9, 10] species distribution models.

Regression-based methods are mainly data-driven and correlate the observed species distribution with different covariates, such as environmental variables and/or those describing human activities. The advantage of these methods is that they generate results mainly driven by empirical data, without strong reliance on hypothetical mechanistic behavioural assumptions. In addition, regression methods can consider spatial or temporal patterns that are not explained by the considered covariables (i.e., phenomenological terms, such as trends), which is important for modelling realistic spatio-temporal changes. A possible drawback of regression models, however, is that they provide information about correlations but not about causalities, which must be considered when interpreting the results. For example, a detected correlation between seabird abundance and fishing vessel density may indicate that the birds are attracted by the fishery process itself (e.g., due to discarding [11]), or, in contrast, they could be disturbed by fishing vessels, but the birds and ships may both be attracted by a locally high fish abundance, eventually leading to a pseudo-correlation of both variables (”third-variable problem” [12]). Furthermore, the explicit nature of regression approaches (i.e., the direct correlation of animal abundance with explanatory variables) limits the complexity of the processes that can be detected/described. In particular, if spatial redistribution processes are considered, implicit equations (e.g., comprising derivatives of the variables of interest) are frequently required to detect (and predict) such processes that explicit equations (e.g., described via regression equations) cannot adequately describe. Finally, predictions based on regression approaches usually work well for parameter values within the experimentally assessed range (i.e., for interpolations), but predictions based on covariables outside the experimental range (i.e., extrapolations) are frequently connected to high uncertainties, and unrealistic results can be obtained.

In contrast, mechanistic modelling approaches (also called “process-based models”) are strongly based on *a priori* (e.g., independently derived) information about species behaviour, and are thus based on specific mechanistic assumptions about the process of interest. The advantage of these approaches is that these assumptions usually constrain the simulations such that predictions (particularly extrapolations) are more robust than in regression approaches [9, 10]. The challenge of these models, however, is that details of specific behavioural parameters are often not known (or their transferability from other studies is questioned), so that simulations/predictions frequently contain speculative elements. Partial differential equations (PDEs) are a class of mechanistic models that can describe various spatial ecological phenomena, and which are particularly suited to describe various types of animal movement and redistribution [13, 14]. In contrast to individual-based mechanistic models (e.g., [15, 16]), PDE-based approaches are not limited to a maximal considered system size (e.g., a maximal number of interacting individuals due to computational constraints), and simulated processes consider the population rather than the individual scale, which makes them particularly suitable for considering numbers of individuals in large study areas, and also for combining with animal density-based regression methods, which also adopt a population-based rather than an individual-based point of view. Despite these factors, PDE-based models are rarely applied in an ecological context, possibly because their implicit nature (i.e., the use of spatial and temporal derivatives) requires a relatively deep mathematical understanding, both for setting up the models and for carrying out simulations. Their implicit nature, however, makes it possible to describe more complex processes, such as spatio-temporal animal redistribution, which cannot be adequately described (i.e., detected and predicted) via explicit correlations alone.

A prime example for studying the impact of human activities on wild animal populations is provided by different marine offshore habitats, where human activities (such as offshore windfarms (OWFs)) are currently increasing, distinctly changing animal distributions and habitat properties [17, 18]. Various modern regression-based approaches have recently been presented and applied to estimate the effects of OWFs on offshore seabird distributions and numbers (e.g., [17, 18, 19, 20, 21]). These methods adequately describe local losses or gains in bird densities due to human pressures (e.g., in the 20 km surroundings of OWFs) and can also detect regional trends caused by several interacting drivers; however, they cannot adequately detect and describe (number-conserving) large-scale movements/displacement, which is required in addition to local losses/gains to assess and predict the effect of regional human activities on trends, animal numbers, and distribution patterns (Fig. 1). Thus, the above-mentioned approaches usually neglect number-conserving spatial animal reorganisation (such as large scale displacement, see Fig. 1 lower right) on the macro-scale, and only measure changes (such as losses) in the surroundings of OWFs (see Fig. 1, upper right). Here, impact-assessments are either based on the interpretation of regional trends or estimated losses in the OWF surroundings, both potentially leading to biased results regarding the information on the large scale reorganisation and potential region-wide abundance changes and trends. OWF impact studies for similar species and regions (e.g., loons in German North Sea waters) thus estimated strongly comparable losses in the OWF surroundings and similar trends, but came to quite different conclusions, because it was difficult or impossible to estimate the contribution of OWFs to the measured regional trends (e.g., [22, 23, 18]). Here (and in the following), “population” does not necessarily refer to a spatially closed population, but rather concerns spatio-temporally varying (possibly migratory) animals intersecting with a specific area of interest – in our example the German exclusive economic zone (EEZ) within the North Sea.

The above advantages and shortcomings of correlative and mechanistic approaches in the context of impact studies suggest the need to either apply both methods in a comparative manner, or alternatively to develop integrated (hybrid) methods to clarify and predict spatio-temporal species distribution and population trends under changing conditions (e.g., [9, 24, 25, 26, 27]). In this study, we followed the latter strategy to improve the estimation and prediction of the changes in animal large-scale distribution and numbers caused by regional human activities, by synergistically joining mechanistic and regression approaches. Notably, we extended previous regression approaches by making additional use of the predictive capacity of mechanistic PDEs, especially when measuring and predicting complex animal redistribution processes beyond simple correlations. We exemplarily applied our approach to investigate the current and future impacts of intensified OWF implementation on common murres (*Uria aalge*) in the German North Sea.

## Materials and Methods

### Study area and data

#### Study area

The exemplarily investigated study area is given by the German EEZ and the offshore parts of the German territorial waters (in the following abbreviated to EEZ) of the German North Sea (Fig. 2B), where several wind farms have been established within the last two decades (Fig. S1A vs. B) with a capacity of 7.7 GW at the end of 2020. Current German Government plans include the implementation of at least 30 GW of offshore wind power in the German EEZ by 2030, at least 40 GW by 2035, and at least 70 GW by 2045 (The Federal German Government 2021; Deutscher Bundestag 2022). The current German marine spatial plan (MSP) designates priority and reserved areas for OWF implementation as shown in Fig. 2B. The draft of the site development plan, however, defines additional areas for OWF development to reach the implementation of 70 GW by 2045. Due to the ongoing planning process, we focused on the areas outlined in the MSP.

#### Raw bird-count data and data processing

A synopsis of the present approach is given in Fig. 3. Several technical details, e.g., with respect to imperfect animal detection during surveys, spatio-temporal autocorrelation, data pooling, and regression analyses are described briefly below, and a more detailed description can be found in [29]. Raw bird-count data were obtained from aerial (digital- and observer-based) as well as vessel (observer-based) surveys from 2003-2020, covering the main period of OWF construction within the German North Sea. Data were obtained from various seabird monitoring and research projects conducted by the University of Kiel (e.g., the German Marine Biodiversity Monitoring on behalf of the Federal Agency for Nature Conservation) and from environmental impact studies and monitoring during construction and operation of wind farms provided by the Federal Maritime and Hydrographic Agency – BSH. Further details of data sources and field methods are given in e.g., Ref.[17, 19, 22]. The study was restricted to the species-specific autumn period (16.07 – 30.09 [30]), as the season with highest numbers of common murres in the German North Sea. In addition, murres are particularly sensitive in the treated season because they are temporarily unable to fly due to moulting and are also in the final stages of rearing young birds. Several technical details regarding data processing prior to the analysis (e.g., with respect to imperfect detection, spatio- temporal autocorrelation, and data pooling) can be found in e.g., Ref.[17, 19, 20, 29]; in the following, only differences from these approaches are detailed.

**Figure 3.**
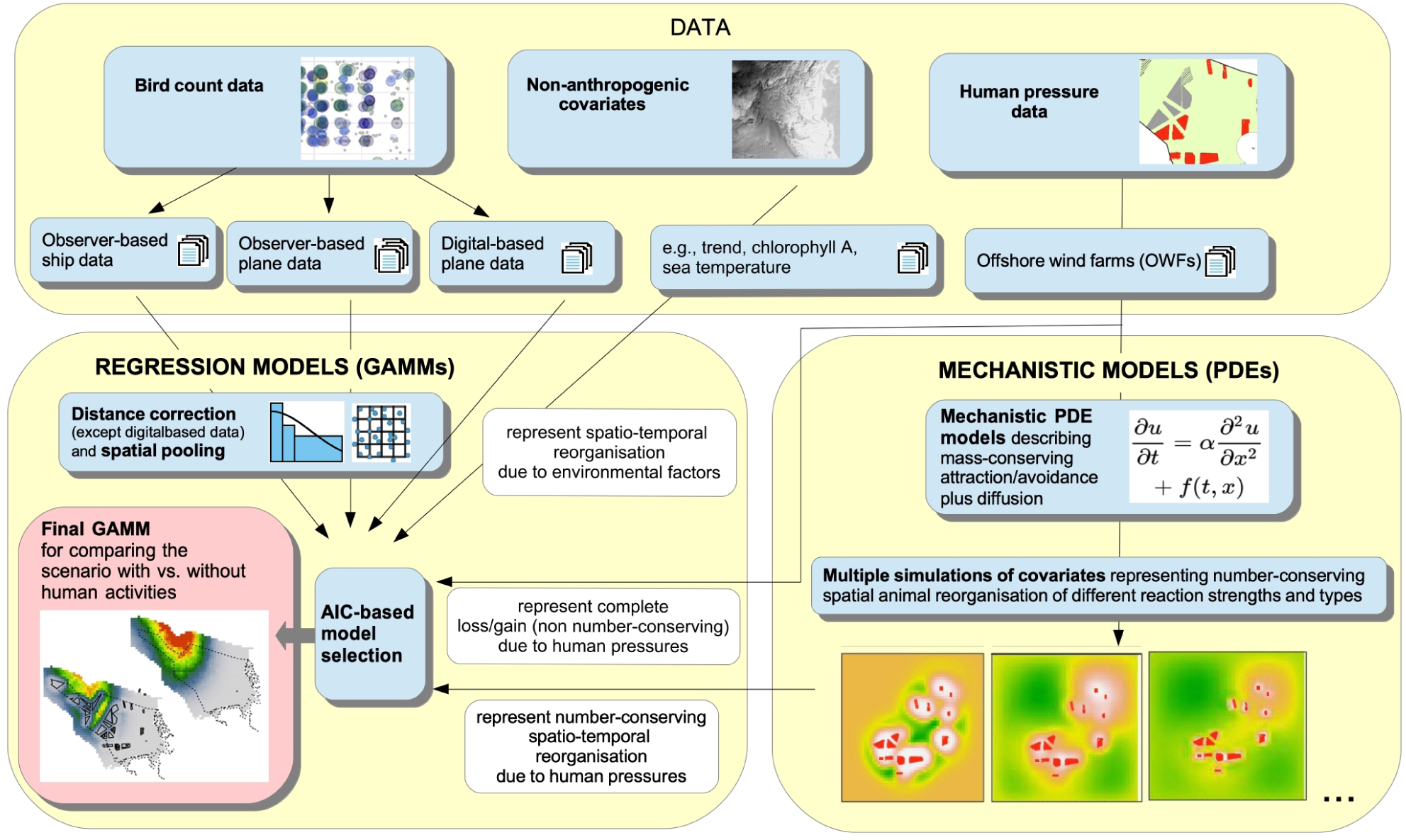
Technical overview of the present approach synergistically combining regression and mechanistic modelling to investigate and predict the effects of regional human activities on animal distribution and numbers within an area of interest. The approach has been exemplarily applied to investigate the impact of intensified OWF activity in the German North Sea on the regional common murre population during autumn.

Distance-corrected bird-count data, locally surveyed area sizes, and covariate values were spatio- temporally pooled using a regular 8 *×* 8 km spatial grid, depicting the optimal compromise between a manageable autocorrelation while keeping a sufficient spatial resolution [20, 29]. In particular, bird-count data and monitored areas were summed separately for each unique combination of grid cell-ID, year, and season, whereas all other covariates (including spatial coordinates) were averaged. This led to a much higher effective spatial resolution than a 8 *×* 8 km than if grid-cell centres had been used instead [29].

Spatial plots of the pooled common murre bird-count data are given in Fig. S1A–B. Final pooled data (without data from OWF construction phases – c.f., below) comprised 9,918 data points with 30,186 observed individuals based on 44,356 monitored km^2^. In particular, 18,480 individuals were counted before construction of the OWFs, whereas 11,706 individuals were counted in phases where OWFs were operating.

#### Data on human activities

Data on wind turbine locations from 2003-2019 were obtained from OSPAR ODIMS (link), and construction- and operation-phase dates for several OWFs were acquired from various online sources. In addition to OWFs within the German EEZ, those close to but outside the EEZ boundary (namely those belonging to the Danish Horns Rev complex and the Netherlands Gemini parks) have also been considered. Planned OWF areas (priority and reserved areas) for future implementation were obtained from the MSP for the German EEZ 2021 (link). Before merging pooled bird-count data with spatio-temporal OWF information, we removed data from the OWF-dependent construction phases to reduce the complexity of the analysis. Subsequently, the distance to the nearest operating OWF was assigned for each (pooled) spatio-temporal observation (with respect to the time point of the observation), with distances *>* 20 km set to 20 km (which implies that no effect is measurable above 20 km [17, 19, 18] and therefore all values set to 20 km are assumed to be unaffected with respect to the meso-scale). Recent results suggest however that the effects at the meso-scale may extend somewhat further than 20 km [31], so our approach may present a (slightly) conservative approach on this point. However, since large-scale effects are captured by the mechanistic components in our model, we assume the corresponding bias as negligible.

#### Environmental covariates

We also considered the effect of natural covariates on birds by investigating different variables potentially representing prey availability and/or suitable habitats. Annual spatial chlorophyll A patterns were obtained from the OceanColor Web provided by NASA (link), as a proxy for annually varying primary production. In addition, spatially varying sea surface temperature was obtained from the NASA MODIS database (link). To also detect and describe oceanographic fronts, which are frequently sites of enhanced physical and biological activity potentially attractive for seabirds [32], we also calculated spatial gradients of sea surface temperature as covariates (which is however only possible for rather static fronts).

After careful analysis, we excluded the additional variables “water depth” and “nearest distance to the coast” from the analysis, because these measures appeared to be collinear with OWF locations, thus strongly confounding the impact study and inflating final standard errors. This bias effect could be eliminated by applying a before-after control-impact (BACI) design (e.g., Ref. [17, 19, 33, 31]); however, a BACI-design was not appropriate for the present study, which focused on a realistic description of time-dependent bird reorganisation in the entire study area, rather than estimating unbiased regional (OWF-specific) reduction effects by introducing two distinct periods (‘before’ vs. ‘after’). Nevertheless, as described below, it was possible to estimate and integrate values from a BACI approach to correct for potential corresponding bias (the latter resulting from scenarios where bird densities correlated with locations of future OWFs – e.g., due to preferences with respect to water depth or distance to the coast). To also consider spatially inhomogeneous bird patterns not explained by the above predictors, we additionally used the spatial coordinates as smooth 2D predictors restricting the maximal number of knots to nine (preventing collinearity between the spatial smooth and OWF locations).

### Mechanistic PDE models and simulations

Our mechanistic model of animal redistribution is based on the Keller-Segel model, originally developed to describe the interplay between directed chemotaxis and random (diffusive) movement of biological units (such as bacteria) [34]. Here, the density *n*(*t*; *x*) of cells and the chemoattractant concentration *c*(*t*; *x*) were considered, where *t* represents the time and *x* a 2D space variable. The interplay of chemotaxis in combination with the production of the chemoattractant is given by the following system of two coupled PDEs, namely:

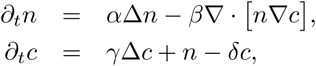

for *t >* 0, *x ∈* R^2^ and appropriate initial and boundary conditions. Here, *∂_t_* depicts the time derivative, Δ the Laplace-operator, *∇* the spatial gradient, and *∇·* the divergence operator. In the first equation, Δ describes diffusive/random (isotrop) movement behaviour scaled by the factor *α*, whereas the term ∇ · [*n∇c*] describes the directed attraction movement along the gradient of *c*. The second equation describes the secretion of *c* by the biological units *n*, including diffusion and degradation.

We adapted this model to describe animal displacement induced by regional human activities. First, in our model, *n*(*x⃗*; *y*) describes the density of animals (with year *y* and spatial location *x⃗* = (*lon*; *lat*)). The variable *c*(*x⃗*; *y*) represents the spatial impact range of a human pressure of interest (c.f., below), whereas the location of the latter is given by the variable *ζ*(*x⃗*; *y*). In our exemplarily application, *ζ*(*x⃗*; *y*) describes the location of OWFs as a binary variable (inside vs. outside the OWF). *ζ*(*x⃗*; *y*), however, is not used directly but is pre-processed to incorporate the fact that the impact is usually not limited to the location of the anthropogenic structure, but a long-range (attraction or avoidance) effect on animals may be exerted, e.g., induced by visual perception, spatial memory [35], sound, or vibration [36]. Indeed, for OWFs, it has been demonstrated that seabirds can be impacted over distances beyond 20 km [22, 17, 18]. Thus instead of using *ζ* directly, we considered the solution *c* of the diffusion equation, *∂_t_c* = *γ*Δ*c* with initial conditions *c*(*x⃗*; *t* = 0) = *ζ*(*x⃗*; *y*), evaluated separately (after 8 numerical time steps *δt* – a heuristically determined value) for each year *y*. This introduces a ‘halo’ around the human activity (exponentially decaying with distance), scaled by the parameter *γ*. The resulting spatio-temporal variable *c* is subsequently rescaled to the range [0, 1] across all considered years *y*. *c*(*x⃗*; *y*) thus represents the spatio-temporally varying human pressure *ζ*(*x⃗*; *y*), augmented by an isotropic long-range effect with strength *γ*. In particular, for *γ* = 0, *c* coincides with *ζ*, whereas for an increasing *γ*, *c* spreads increasingly from the original source of human activity. Importantly, this approach naturally also incorporates the effect that larger OWF areas induce a stronger effect than small ones, the latter being easily “smoothed out” by the applied diffusion. Thus final *c*-values for larger compared with smaller OWF areas are larger in both extent and absolute (maximal) value. Examples of two spatial patterns of *c* for different values of *γ* are given in Fig. S2, considering operating OWFs in German North Sea waters in 2019.

Thus, for each year *y* separately, we first considered (and numerically solved) the following PDE describing the human-induced pressure by:

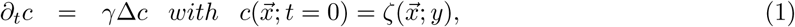

and subsequently the PDE simulating animal redistribution by:

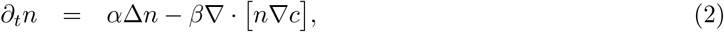

both for a defined number of time steps *δt* (c.f., below). Here, *∂_t_* is the time derivative with respect to the numerical time variable *t*, *γ* scales the spatial reach of the impact of the human activity, *α* quantifies the amount of undirected (random-like) animal movement, and *β* scales the strength of avoidance or attraction. Examples of abundance patterns resulting from different strengths of *β* are given in Fig. S2. Numerical simulations after eight numerical time steps *δt* were considered separately for each year. As initial conditions in our simulations, for *n*(*x⃗*; *y*), we used the constant value of *n*(*x⃗*; *y_start_*) = 1 for the first considered year 2003, and the respective foregoing solution *n*(*x⃗*; *y −* 1) for all following years. Thus, if human activities changed in time, the stepwise spatial reorganisation of animals due to the changing pressure locations and strengths was simulated. Furthermore, we used Dirichlet boundary conditions with a constant abundance of *n*(*--x_bound_*; *y*) = 1, representing an unchanged population density at the boundaries of the considered square. The boundaries of this square (longitude extent 3.0*^◦^*, 9.0*^◦^* and latitude extent 53.0*^◦^*, 56.5*^◦^*) were either close to the coastline, or sufficiently far away from the boundaries of the considered study area that their influence on abundance patterns with the study area could be neglected.

A regular spatial grid with 12,100 grid cells was used during PDE simulations. The final annual and spatially given variable *n* was logarithmised (due to the use of a log-link function in regression-based approaches – c.f., below) and rescaled to the range [0, 1] across all considered years *y* (for better convergence properties during regression analyses), eventually leading to the variable *sim OWF* used as a spatio-temporal predictor of large-scale animal distribution with respect to the OWF location during regression-based modelling (c.f., below).

In summary, the variable *sim OWF* is proportional to animal density and represents spatial macro-scale reorganisation (especially active, individual number-preserving movements in space – cf., Fig. 1) of the birds in response to the implementation of the OWF. It is given by the solution of differential equations originally developed to describe chemotaxis and can be used in the following as additional predictor in regression-based species distribution modelling.

### Integrative generalised additive mixed model (GAMM)-based regression modelling

The scheme for synergistically joining PDE- and regression-based methods is shown in Fig. 3 and is given by a two-step procedure. In the first step, several thousand different possible anthropogenically induced redistribution patterns were simulated based on mechanistic PDE models (c.f., above) and subsequently used as potential predictors in regression GAMM-based species distribution models, including the simultaneous consideration of additional (natural and/or phenomenological) predictors. In the second step, model-selection procedures were applied to select the simulated pattern that best explained the empirical animal redistribution data, eventually leading to the final regression model used for the impact analysis.

The detailed scheme is described in the following. First, 1,000 different PDE-based simulations of animal redistribution (Eq. (1)-(2)) were performed, where each simulation used stochastic values for *γ ∈* [0, 100.0], *α ∈* [0, 0.1], and *β ∈* [0, 100.0] (for an explanation of these parameters, see the derivation of the mechanistic formulas and/or Fig. S2). Reasonable ranges for these parameters were determined heuristically. GAMMs [37, 38, 39] were applied separately for each run, using the corresponding simulated pattern (variable *sim OWF* – c.f., above) as predictor, along with other investigated covariates. In particular, the most comprehensive GAMM was given by:

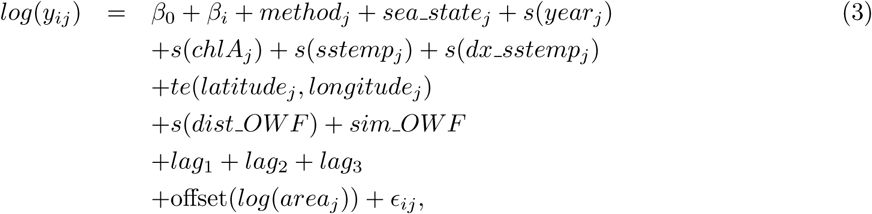

with *E_ij_ ∼ N* (0*, σ*_1_^2^) and *β_i_ ∼ N* (0*, σ*_2_^2^) normally and independently distributed. Here, *y_ij_* is the vector of (pooled, distance-corrected) bird numbers, where the index *j* refers to the sampling unit number and the index *i* to the random intercept representing intra-seasonal changes by defining intervals of 10 days. *β*_0_ is the fixed intercept, *s*(.) depicts a smooth regression spline (with the optimal number of knots estimated via generalised cross-validation), and *te*(.) a tensor product 2D spline [38]. *method_j_* and *sea state_j_*are related to the observation platform and wind conditions, respectively, both influencing the distance- independent detectability (both variables are however also considered in an additional distance-dependent detection correction step [29]). *s*(*year_j_*) represents a (possibly nonlinear) trend, frequently influenced by various factors intrinsic and extrinsic to the study area (c.f., above), and *s*(*chlA_j_*) represents the (possibly nonlinear) dependency on chlorophyll A concentration. *sstemp_j_* is the sea surface temperature and *dx sstemp_j_* is its spatial gradient. *latitude_j_* and *longitude_j_* are spatial coordinates, and the corresponding 2D smooth is restricted to maximal 9 degrees of freedom (heuristically determined), as described above, to avoid collinearity. *s*(*dist OWF*) measures a local (possibly nonlinear) bird density change with distance to the closest operating OWF, whereas *sim OWF* represents the simulated PDE-based large-scale bird redistribution due to OWFs (c.f., foregoing section). In particular, *sim OWF* captures OWF-driven number-conserving spatial reorganisation of animals on spatial macro-scales, whereas *s*(*dist OWF*) measures the loss (or gain) of animals at the meso-scale (kilometres) surroundings of the OWFs, which cannot be explained by macro-scale movements but rather represent a complete loss (or gain) of animals from the considered area. Notably, both OWF-related variables might also reflect positive/attraction effects, in which case, a negative correlation with these variables would be estimated during regression. Because survey effort varied per sampling unit, the logarithm of the locally surveyed area was included as an offset [40]. The terms *lag*_1_*, lag*_2_*, lag*_3_ finally refer to potential spatio-temporal autocorrelation. For more technical details, see [29].

The appropriate probability distribution with respect to the subsequently applied GAMM was first selected for each of the 1,000 PDE-based simulated patterns separately (based on the Akaike information criterion (AIC) [41]) from the negative binomial [42] and the Tweedie distribution [43], using the above presented most comprehensive model. Next, using this preferred probability distribution, an AIC-based predictor selection was applied, testing 42 biologically reasonable subsets of the predictors *s*(*chlA_j_*), *s*(*sstemp_j_*), *s*(*dx sstemp_j_*), *s*(*dist OWF*) and *sim OWF*, where *log*(1 + *dist OWF*) instead of *dist OWF* was also tested, to better describe meso-scale effects / “footprints”[20]. The model with the lowest AIC value was finally selected. To exclude possible unrealistic models *per se*, models in which both OWF-related variables were contradictory (i.e., one variable suggested an attraction while the other measured a disturbance) were excluded; however, this was rarely the case in practice. This finally led to 1,000 different AIC-values (related to the different simulated PDE-based patterns), importantly including those models in which OWF-related variables were not (or only partially) selected. In a second step, the entire simulation procedure was repeated, narrowing *γ*-, *α*-, and *β*-parameter ranges to those associated with the models showing the 100 lowest AIC values from the first run. Thus, the first run evaluated the entire reasonable PDE parameter space, whereas the second run was restricted to the region where models showed low AIC values, thus locally refining the global AIC minimum, the latter representing the most appropriate simulated PDE-based pattern describing OWF-induced spatial animal reorganisation.

In the last step, the final species distribution model (termed “sdGAMM”) was defined and fitted and subsequently used to analyse the current and future impacts of selected human activities (c.f., below). Here, to increase the robustness of the PDE-based results (respectively to minimise the selection/influence of random correlations due to the high number of tested predictors), average PDE parameters from the five models with the lowest AIC values and comprising the predictor *sim OWF* were used to generate the final variable *sim OWF* based on the PDE scheme described above. Using this final variable, predictor selection (c.f., above) was again applied to obtain the final sdGAMM. Model validation was done using various residual-based plots, e.g., as described in [40, 44].

### Population estimates, predictions, and uncertainties

The geographical square used as the virtual area for PDE simulations (c.f., above) was used as a basis for model-based predictions. This area was further restricted to the German EEZ augmented by a regular belt of 40 *km* such that the total considered area (concerning water-covered areas) roughly corresponded to twice the area of the German EEZ. The rationale behind this restriction was that the belt lying outside the study area contained sufficient area to incorporate displaced birds, while the geographic extrapolation was distinctly limited, preventing unrealistic predictions (e.g., with respect to the estimated preference of environmental covariates). Thus, the expression “number conserving” in this work concerns the region including the EEZ as well as the above-mentioned belt. However, assuming that this belt was an undisturbed area that might take up displaced birds was only true to a very limited extent when considering future scenarios, due to OWF plans by neighbouring countries not included in this study.

This spatial prediction data frame was subsequently multiplied by and assigned with the different years considered in this study, from 2003–2020, in a first step by merging the yearly spatial data with corresponding spatio-temporal information from all the above considered covariates/predictors. For future predictions, 10 additional “virtual” years were then augmented in a similar way, using the values from 2020 for all covariates, except the two OWF-related variables. To investigate the impact of planned OWF extensions in German North Sea waters, we first simulated the implementation of “priority areas” (grey-shaded areas in Fig. 2B) in virtual years 1-5 (including the corresponding calculation of the variables *dist OWF* and *sim OWF*), and the “reserved areas” (grey-striped areas in Fig. 2B) were similarly integrated for the subsequent 5 virtual years. Notably, the exact number and time points of years used here were rather arbitrary, with minor impact on the predictions, given that predicted/simulated bird patterns appeared to stabilise quickly after each virtual OWF extension (c.f., Fig. 4A–B).

**Figure 4.**
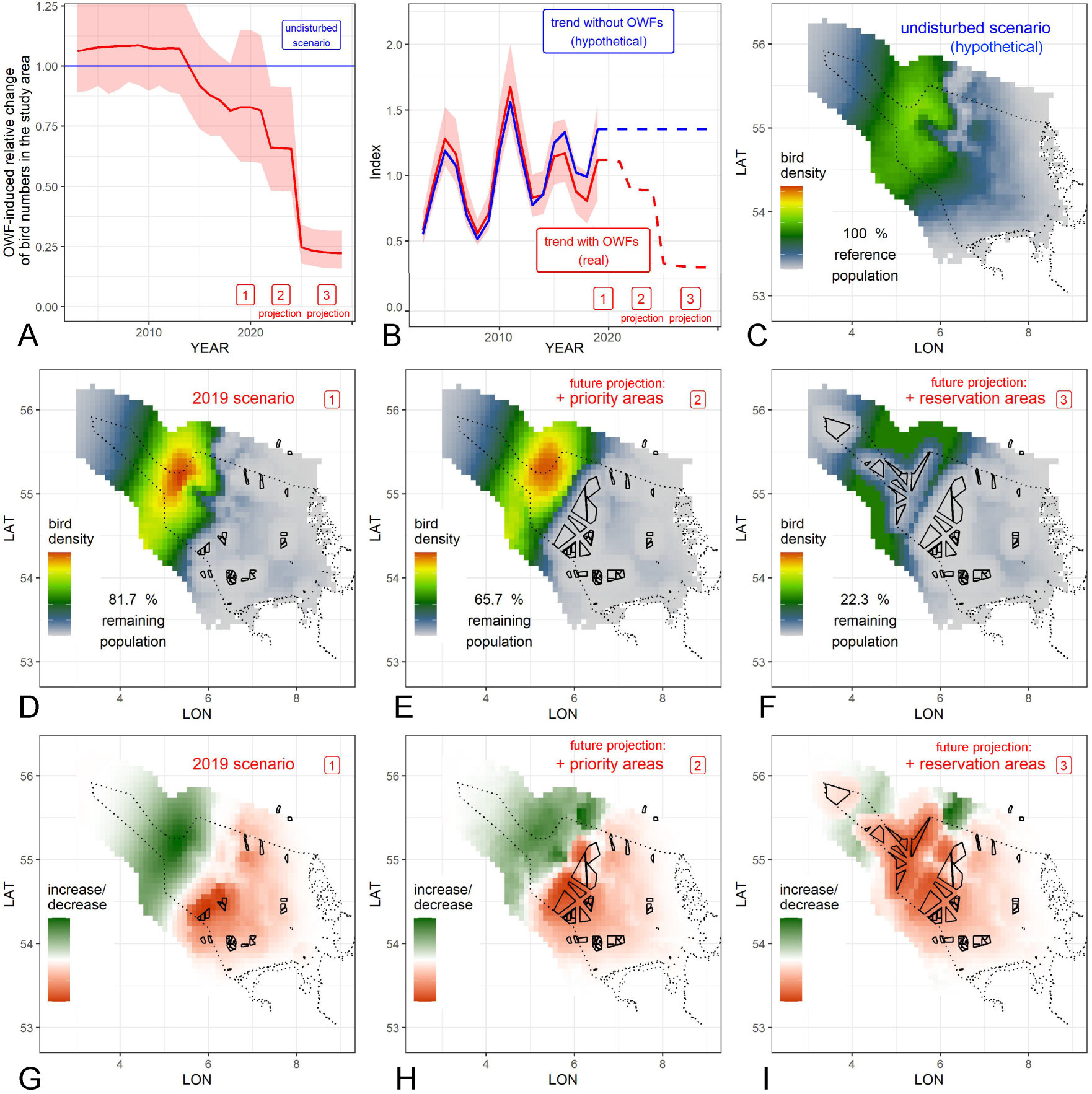
Predicted changes in numbers and distribution patterns of common murres during autumn in the study area (German EEZ and coastal waters) and surroundings. (A) Relative changes in bird numbers within the study area (solid red line) due to OWFs compared with predicted number without OWFs (blue line). Red-shaded areas: 95% confidence bands. (B) Nonlinear trend estimates (trend index rescaled to a mean of 1.0) including the effect of existing (solid red line) and planned (dashed red line) OWFs. No OWF-independent trend is assumed from 2019 onwards, due to difficulties of extrapolations. Blue line: analogous but hypothetical scenario without OWFs. (C) Reference spatial bird distribution pattern considering the hypothetical scenario without OWFs. (D)–(F) Spatial bird densities with different OWF scenarios. (G)–(I) Local relative bird density changes due to different OWF scenarios, compared with hypothetical undisturbed scenario. For better visualisation, bird density changes (G–I) were rescaled separately for each plot and separately between -1 and 0 as well as 0 and 1 (i.e., with respect to increase (green) vs. decrease (red)), and the decrease and increase strengths are thus not directly comparable. In contrast, the colour scaling in (C–F) is directly comparable. All OWF scenarios later than 2019 are fictitious (but are based on currently discussed/planned scenarios in the German EEZ).

The final sdGAMM (as derived above) was used as a basis to predict bird densities in real and hypothetical scenarios. In particular, the estimation of anthropogenically induced changes was mainly based on comparisons between the (past, present, or future) disturbed scenario (with OWFs) vs. the hypothetical scenario in which no OWFs were constructed. These two scenarios were predicted for each year separately and related to each other as follows: first, the situation comprising existing, present, or future OWFs was predicted, using similar predictor variables to those used during the original fit of the model (possibly augmented by planned OWF data if future scenarios were considered – as described above). Second, the same data were used setting the variable *dist OWF* constant to 20 *km* (representing only areas outside the meso-scale impact of OWFs) and also setting *sim OWF* to its initial constant value (i.e., no OWFs driving macro-scale spatial reorganisation of birds) – thus, the hypothetical scenario without OWF disturbance is considered.

As described above, the local effects of OWFs on bird distribution were captured by the variable *dist OWF*, which measures the distance to the closest operating OWF. However, these results may be biased if birds already showed particularly high or low densities in areas with future OWFs before OWF construction, either by chance or by co-localisation of environmental covariates such as depth or distance to the coast. In particular, our previous studies on common murres (e.g., Ref. [31]) showed that a non-BACI approach tended to underestimate the strength of local avoidance for this species (non-BACI results not published), and a BACI design has to be applied to overcome this bias. As noted above, however, this is not possible within the present spatio-temporal framework because it requires the data to be separated into two distinct “before” vs. “after” phases, which is not possible in a temporally consistent way, given that operation phases differ among different OWP clusters [33, 31]. To obtain unbiased estimates of the meso-scale OWF effects, we therefore applied a separate correction step. We first restructured the data according to the BACI-design, as applied in Ref. [31] and fitted the above sdGAMM (including, if selected, the variable *sim OWF*) but replacing the predictor *s*(*dist OWF*) by the expression *OWF* + *PERIOD* + *OWF* : *PERIOD*, where *OWF* is an indicator variable for birds in the 20 km surrounding of existing or future OWFs, and the interaction term *OWF* : *PERIOD* was used to estimate the unbiased OWF-induced reduction effect (for details on this approach see e.g., [33]). This modified sdGAMM was termed baciGAMM, and the resulting BACI-effect size was compared by the overall reduction induced by the *s*(*dist OWF*) in our regular sdGAMM in the 20 km surrounding of existing OWFs, and a correction factor was finally derived and applied by the quotient of these two measures.

Although PDE-based simulations are in principle animal-number-conserving, the *sim OWF* -induced changes might induce slight violations of number conservation between both predicted scenarios (concerning the German EEZ and a spatial buffer – cf., above). This is partly because preferences for additional (environmental) variables might induce a spatial re-weighting (some places are more preferred than others), which interplays with spatial changes in *sim OWF*. To guarantee bird-number conservation during *sim OWF* -induced changes, we introduced an additional correction step. Here, the disturbed and undisturbed scenarios were predicted separately for each year, but neglecting the influence of *dist OWF* by setting it to 20 *km* everywhere for both predictions (since this variable would induce ‘legal’ changes in animal numbers, which would however bias the correction term derived in this step). Subsequently, the predictions of the undisturbed scenario were multiplied by the factor *N_disturbed_/N_undistrubed_*(where *N* is the predicted total bird number in the entire considered area), which slightly corrects for any number-conserving violations due to large-scale bird movements. In a second step, the prediction of the disturbed scenario was repeated setting *dist OWF* to the values related to existing or future OWFs. Figuratively speaking, the correction step re-weighted the *sim OWF* -induced spatial reorganisation in the hypothetical undisturbed scenario by the local preference with respect to environmental variables (the latter cannot be considered directly during PDE simulations, because only OWF-based variables were integrated to limit the complexity); an approach which we assume will provide the most realistic description of reorganisation – namely a combined influence from OWF-induced replacement and preference of local habitats.

An additional correction step was applied to prevent bias due to extrapolation outside the study area, which may present a potential problem during regression-based approaches (c.f., Introduction section): for each year and prediction separately, maximal predicted bird densities outside the study area (where empirical data were sparse) were locally cropped to the maximal predicted density value inside the study area. In particular, this inhibited unrealistically high predicted densities due to extrapolations outside the empirically assessed geographical range.

Yearly OWF-driven relative changes in bird numbers within the entire study area were finally calculated by dividing the predicted numbers for the disturbed scenario by those of the (hypothetical) undisturbed scenario. Confidence intervals (95%) of these yearly values were calculated based on appropriate quantiles from 1,000 resamples from the posterior distribution of the parameters of the final sdGAMM, in conjunction with the above predictions, i.e., based on the variance-covariance matrix of model parameters [38].

To identify areas with particularly strong declines or increases in bird density due to OWFs, from the spatial distribution pattern of the disturbed scenario (considering a specific year) we pointwise substracted the corresponding values related to the undisturbed scenario. Hence, local absolute, rather than relative changes were evaluated, with local negative values indicating a local OWF-driven decline and positive values indicating an increase. Local decreases were generally stronger and/or more frequent than increases in the considered examples, and positive and negative values were thus rescaled separately between 0 and 1 (respectively -1 and 0) for each considered scenario, to improve visualisation. Hence, in the corresponding subfigures, the intensities of decreases (red) vs. increases (green) are not quantitatively comparable.

### Software

All statistical analyses, validation procedures, and visualizations were performed using the statistical software **R** [45]. All PDE-based analyses were based on the package *ReacTran* [46]. Spatial analyses (amongst others of spatial autocorrelation) were carried out using the packages *sp* [47], *gstat* [48], *rgdal* [49], geosphere [50], *prevR* [51], and *rgeos* [52]. *ggplot2* [53] was used for all other visualisations and plots. Finally, the package *mgcv* [38] was used for GAMM analyses. All computations were performed using a desktop computer with eight physical cores (Intel Core i7–10700K CPU with 3.80 GHz). The computation time for all presented PDE- and GAMM-based analyses (including resampling) was approximately 48 hours (using parallel computing).

## Results and discussion

In this section, we present and discuss the exemplarily evaluated qualitative and quantitative changes in the occurrence of common murres in the German EEZ of the North Sea (including coastal regions), driven by previous and planned OWF implementation within German waters. The final sdGAMM revealed that, for the considered species, both OWF-related variables (*s*(*dist OWF*) and *sim OWF*) were selected as predictors and both appeared to be highly significant (*p <* 0.0001). In particular, both variables showed a distinct negative effect of OWFs on the birds. This observation is in accordance with previous results regarding meso- and macro-scale OWF effects on this species in the German North Sea [54, 31]. Removing each of these variables separately from the sdGAMM led to an AIC-increase of 92.5 AIC units for the variable *sim OWF* and to an increase of 249.4 AIC units for the variable *s*(*dist OWF*), demonstrating that both variables strongly contributed to an improved description of the empirical spatio-temporal bird distribution patterns [55]. The fact that *s*(*dist OWF*) resulted in a significantly stronger change in AIC than *sim OWF* suggests that the effects on bird densities within 20 km of OWFs are more pronounced than the large-scale spatial reorganization. We thus demonstrated two qualitatively different reactions of common murres to operating OWFs: a macro-scale spatial reorganisation, as well as a local total loss of birds within the meso-scale surroundings of the OWFs.

Based on these relationships, we further evaluated and quantified the influence of existing and planned OWFs on common murre population sizes and distribution in the German EEZ, using the predictions of the sdGAMM. Fig. 4A shows the predicted relative change in the German North Sea population size (solid red line) compared with the hypothetical undisturbed scenario (blue line) depending on the year (x-axis) for the different OWF scenarios (red numbered boxes). Notably, a slight (non-significant) OWF-induced increase in common murres in German waters was predicted until 2013. This might be explained by birds being displaced from Danish into German waters due to the early construction of OWFs (the Danish OWF “Horns Rev 1” north of the German AWZ started operation in 2002). However, the effect was non-significant (confidence bands strongly overlap with the blue line in Fig. 4A) and the results should therefore not be over-interpreted. From 2014 onwards, however, our predictions suggested a distinct negative impact on the German common murre population (Fig. 4A), coincident with intensified expansion of OWFs in the German EEZ, with each of the considered OWF extension scenarios causing a distinct step-like decline, which were highly significant for the priority areas and the priority plus reservation areas (confidence bands in Fig. 4A do not intersect the blue line). Here, the drop was steepest for the reservation areas, in accordance with the particular importance of offshore areas for this species, in combination with the fact that adding the reservation areas meant there was no longer a large retreat area available for this species in the German EEZ.

Based on the sdGAMM, we also estimated a nonlinear trend in common murre numbers summed up for the German EEZ (rescaled to an index of mean value 1.0), separately for the scenario with (red line) and without (blue line) existing and planned OWFs (Fig. 4B). Although yearly numbers fluctuated strongly, possibly influenced by large-scale spatial variability across geopolitical borders within the North Sea, OWF-induced predicted losses in priority plus reservation areas led to reductions distinctly below observed natural minima. Importantly, the trends in Fig 4A and B from 2020 onwards are fictitious in that murre numbers summed up for the undisturbed scenario from 2020 onwards are set to a constant value (because influences not resulting from OWFs cannot be reliably predicted) and because the years of stepwise OWF expansion probably do not coincide with the times of actual expansion. However, the latter is irrelevant in the context of these analyses, given that the main focus was on the qualitative and quantitative effects of the construction projects, rather than on exact dates.

The associated predicted spatial changes are given in detail in Fig. 4C–I. Compared with the undisturbed scenario (Fig. 4C), the situation in 2019 already showed a distinct displacement of birds, which accumulated in more offshore regions (Fig. 4D – also visible in the raw data in Fig. S1A vs. B) and leading to a (barely non-significant) loss of 18.3% of the murre population in German waters (i.e., 81.7% remain). In particular, the south-easterly (more nearshore) region, including OWFs, showed large-scale decreases in bird densities, whereas an increase was observed in the north-western (far offshore) part of the EEZ (Fig. 4D,G). This process was intensified by the addition of priority areas (total significant loss in the EEZ: 34.3% – Fig. 4E,H – i.e., 65.7% remain), and in the case of additional OWF expansion within the reservation areas, with nearly the entire EEZ affected by significant losses reducing the population by 77.7% to only 22.3% of its size in the OWF-undisturbed scenario (Fig. 4F,I). Here, visible bird densities only persisted between wind farm clusters and with maximal possible distances to OWFs. In reality, however, even their whereabouts are highly questionable, given that other human activities and thus disturbances, such as shipping, will be concentrated in the remaining areas. In addition, there may also be increased intraspecific competition (and possibly even interspecific competition if other species are also pushed into these areas). From this point of view, our results likely represent a conservative (probably underestimated) prediction of losses (c.f., detailed discussion below).

Our qualitative and quantitative results were within the orders of magnitude predicted in Ref. [31], although this previous study only considered meso-scale effects and not additional large-scale reorganisation. As a consequence, Ref. [31] did not describe the total loss of animals in the German EEZ, but rather the proportion that will suffer habitat loss around the OWF (meso-scale) according to a forecast based on animals being virtually “cut out” based on measured displacement caused by operating OWF. In contrast, birds in the present study could also lose their habitat due to meso-scale effects, but our modelling approach could quantify those that disappeared completely from the study vs. those that flew elsewhere and were therefore not lost to the EEZ. Thus for the priority scenario, the current study predicted a 34.3% loss of animals in the German EEZ (instead of 48% habitat loss in Ref. [31]), whereas for the priority plus reservation area scenario, we predicted a 77.7% loss in the EEZ (instead of 69% habitat loss in Ref. [31]).

The first-mentioned difference between the percentages of both approaches thus reflects the fact that in the 2019 and the first scenario birds accumulate in more-offshore regions of the German EEZ, partly damping the effect of meso-scale habitat loss (i, e., in the OWF surroundings) since not all birds are lost from the EEZ but some just spatially reorganise such that they keep a sufficiently large distance to OWFs but still remain within the EEZ. This difference between the estimates of the two methods is reversed for the second expansion scenario including priority plus reservation areas, since here, the large-scale reorganisation (expressed by *sim OWF*) adds to the losses on the meso-scale, as birds are now pushed beyond the EEZ-boundaries because there are no longer enough undisturbed areas in the EEZ (which might in reality be only present to a very limited extent, since OWFs from other countries might be installed here). Neglecting the large-scale reorganisation can thus lead to both an over- and underestimation of the effects on regional population sizes based on habitat loss only, depending on if and how displaced birds accumulate within and outside the considered area. When comparing the estimated losses of the present study with the recent estimates of Ref. [31], however, various additional factors might result in additional deviations.

Apart from the key difference that the present study augmented large-scale reorganisation to meso-scale losses (and in addition, EEZ population changes relative to the hypothetical undisturbed scenario have been quantified instead of habitat loss), there were also various technical differences in the presented work vs. the study of Ref. [31]: this includes a spatio-temporal vs. a BACI-approach, further differences with respect to data structure and evaluation, distinctly different spatial grid sizes: (8 x 8 km vs. 2 x 2 km), different spatio-temporal covariates, and a smooth dependency on the OWF-distance vs. a corresponding binary variable. Any/all of these differences might influence the final estimates. Despite these differences, the estimated meso-scale effects of both studies were highly comparable in terms of quality and quantity (c.f., Fig. S3 and below). Finally, comparing the results of both approaches, corresponding confidence bands should be taken into account.

When fitting the baciGAMM using different radii around OWFs, the obtained orders of magnitude with respect to meso-scale OWF avoidance were similar to those presented in Ref. [31]: e.g., for a 1 km radius we measured a 91.4% reduction (Ref. [31]: 91%), an 80.3% reduction for 5 km (Ref. [31]: 80%), and a loss of 72.9% for a 10 km radius (Ref. [31]: 76%). This demonstrates good robustness of these estimated meso-scale effects, despite the above methodological and technical differences between the studies. Interestingly, removing the variable *sim OWF* from the baciGAMM (i.e., considering only the meso-scale effects) led to very similar orders of magnitude regarding the meso-scale effects (91.6%, 81.4% and 74.2% for 1 km, 5 km and 10 km radii, respectively). This demonstrates that both meso-scale losses (described by *OWF* or *dist OWF*) and large-scale reorganisation (described by *sim OWF*) captured two distinct processes acting at different spatial scales, given that meso-scale effects did not sensitively depend on the variable *sim OWF*. Indeed, further investigation of the selected/estimated parameters and patterns with respect to the PDE-based predictor *sim OWF* (macro-scale spatial reorganisation) revealed that estimated bird displacement acted at a much larger scale than the local effects given by *s*(*dist OWF*). A spatial plot of the variable *sim OWF* for 2019 is given in Fig. S4, demonstrating again the large-scale effects on this species noted in Ref. [31]. Ref. [31] noted a reduction in common murres up to 21 km distance from operating OWFs in autumn (relative to those between 22-35 km distance and to densities before OWF construction). In the present study, we limited the detection of meso-scale effects to 20 km to distinguish between meso- and macro-scale effects, which represents a slightly conservative approach.

The present results provide relatively conservative estimates of the human-induced effects on the distribution and number of animals for various reasons. First, the current approach did not apply any constraints with respect to predicted local animal densities within the study area. In the considered example, this led to bird densities distinctly beyond those predicted for the undisturbed scenario (c.f., Fig. 4C vs. D,E). In undisturbed situations, such high densities might either not occur (because birds avoid very high densities e. g., due to inter- or intraspecfic competition), or they may occur but lead to additional negative effects not considered within our approach. Second, the present example only considered OWFs, but other human pressures, including OWF-associated ship traffic [17] will increase simultaneously, further adding to the direct impact. In addition, cargo ship activity and OWF related ship traffic will increase in those areas not occupied by OWFs, where our predicted remaining bird densities are highest. Third, planning of OWF areas in the German North Sea is currently ongoing, and the recent draft spatial development plan outlines even larger areas for OWF expansion, especially in the priority and reserved areas (BSH 2024). Fourth, our predictions assumed that areas outside the German EEZ could be used unreservedly by birds as evasive areas; however, adjacent countries may have similar OWF extension plans that might restrict evasion and thus make our prediction with respect to bird accumulations in the surroundings of the German EEZ too optimistic.

The predicted declines in abundance of common murres as a result of planned OWF expansion in the German North Sea are dramatic. Especially in combination with the above reasons leading to an underestimation of the effects, it is not unrealistic to consider that this species might almost completely disappear from the German North Sea if OWF expansion is also realized in the reservation areas.

## Conclusions and outlook

In summary, we present a new analysis framework to estimate and predict changes in animal distribution and numbers caused by increased human activities within a specific area of interest. In particular, the framework synergistically uses mechanistic- and regression-based modelling approaches to describe animal redistributions, along with regional losses/gains of animals as a result of human activities. By directly comparing the predicted scenarios with and without human pressures, the present approach does not rely on the problematic interpretation of regional trends, which may be influenced by various intrinsic and extrinsic processes that are difficult to estimate. In addition, this approach can be used to not only understand previous and current human-induced changes, but also to predict the impact of different hypothetical future scenarios.

We exemplarily applied our approach to investigate the current and future impacts of intensified OWF implementation on the numbers and distribution of common murres during autumn, within the German North Sea. We demonstrated that OWFs may have a distinct negative impact on this species by inducing both their redistribution, as well as their loss from the study area. In particular, OWFs constructed up to 2019 reduced common murre numbers in German waters by approx. one fifith. The addition of OWF priority areas (representing planned expansions in the German EEZ up to approximately 2030) increased this reduction to approximately one third. Finally, if the OWF reservation areas are also augmented by optional areas for further future expansions in the German EEZ up to 2045, the predicted German population loss increases up to four fifths. These results are particularly concerning given that these estimates are conservative for several reasons, suggesting that if all planned priority and reservation areas were used for OWF implementation, the population of common murres within the German EEZ may be completely lost. However, it should be emphasized that these shifts and decreases are always based on the current behavior of the common murre. The scenario could therefore change in the event of habituation.

The predictive capacity of the present approach may be used in the near future to estimate and compare the impacts of alternative strategies, e.g., during the expansion of alternative energy technologies in previously less-disturbed habitats. Optimising both spatial and temporal implementation choices will help us to minimise the human impact on wildlife, with the ultimate aim of preserving biodiversity while fighting climate change.

## Acknowledgements and funding

The following environmental consulting companies also recorded bird-distribution data: BioConsult SH GmbH, IBL Umweltplanung GmbH, and Institut für Angewandte Ökosystemforschung GmbH. Major parts of the data collection and analyses were conducted within the research projects OWP-Seevögel (FKZ:3520860100). S. Furness provided linguistic support.

## Author Contributions

SG/VP conceived the ecological aims of the study, and SG led its coordination. MM developed the new approach conceptually, developed the R-codes and computational methods for PDE and GAMM analyses, did the visualisation, and performed all analyses and statistical validations. All authors discussed and developed the required features of the analysis. SG/VP/KB/NM/HS/VD provided biological input and participated in validation of the results in an ecological context. All authors helped to draft the manuscript and read and approved the final manuscript.

## Supporting information

**Figure S1.**
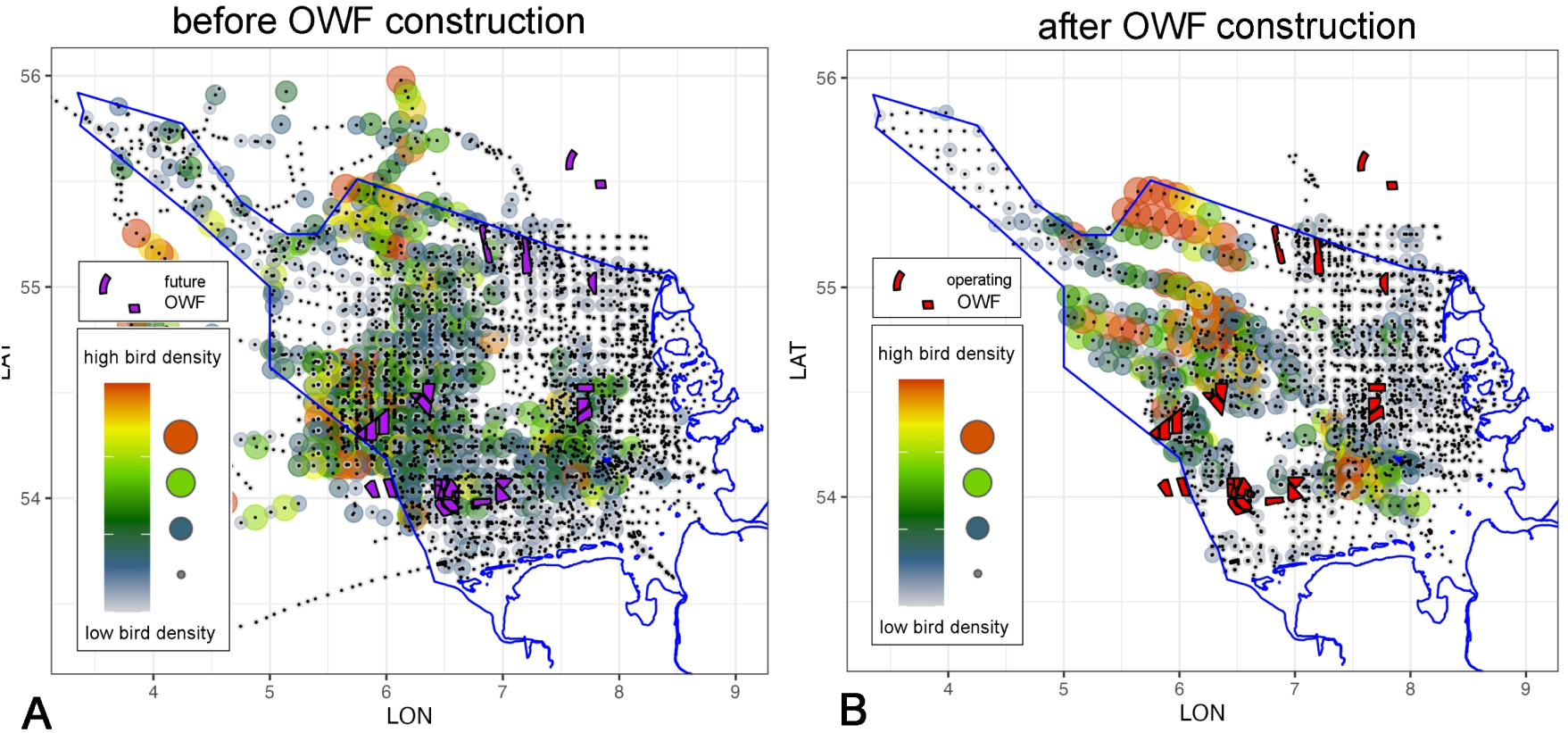
Spatio-temporally pooled raw bird-count data (logarithm of murre number per km^2^) from autumn before OWF construction (A) and after OWF construction (B). Black dots represent all (pooled) observations (zero and non-zero), coloured dots and their size are related to non-zero bird densities, purple areas in (A) indicate areas with future OWFs (before construction) and red areas in (B) represent operating OWFs. Notably, (A) and (B) do not present a temporally homogeneous scenario, because construction and operation phases may differ among OWFs.

**Figure S2.**
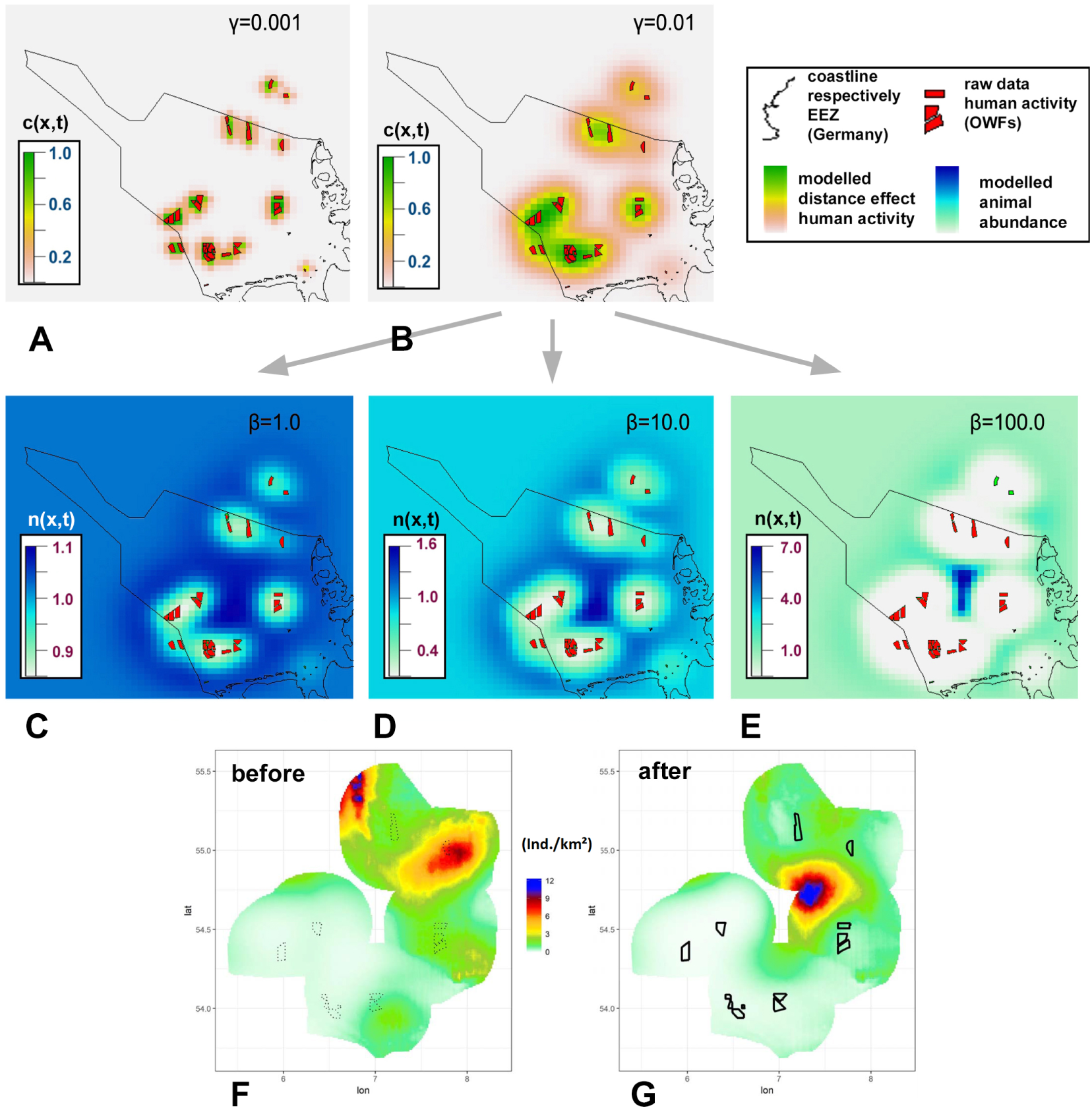
Examples of mechanistic simulations considering the influence of offshore wind farms (OWFs – red) within German North Sea waters in 2019 (black lines represent coastline and boundaries of the exclusive economic zone (EEZ)). (A)-(B) Spatial distribution of the variable *c*(*t*; *x*) representing OWF locations (red) augmented by an isotropic long-range effect with strength *γ* (beige/yellow/green colour range depicts *c*(*x*; *t*)), evaluated for two different values of *γ*. (C)-(E) Example simulations of animal redistribution (avoidance) based on the OWF-related variable *c*(*t*; *x*). Different strengths of animal attraction/avoidance scaled by *β* (subfigures below; colour range represents *n*(*x*; *t*)), leading to quantitatively and qualitatively different animal distribution patterns. (F)-(G) Actual loon distribution before (F) vs. after (G) OWF construction in the German Bight (from Ref. [22]), in the after-period strongly resembling simulated pattern in (E).

**Figure S3.**
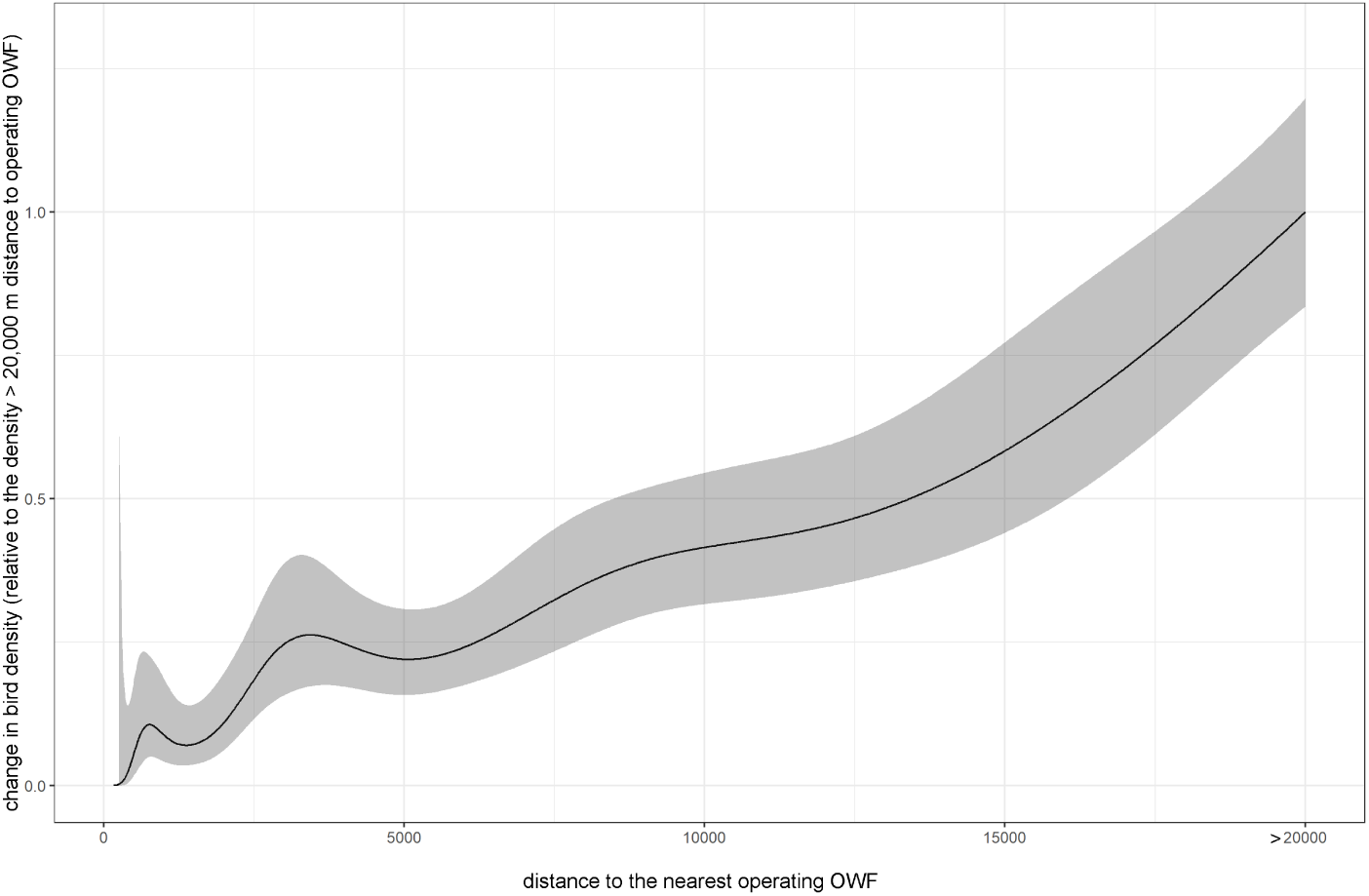
Estimated relative loss in bird density on the meso-scale, i.e., depending on distance to the nearest operating OWF. These meso-efffects are directly derived from the sdGAMM in this study (slightly modified, c.f., below), BACI-related correction factors have not yet been applied (c.f., below). For a better comparability to the results in Ref. [31], the variable *sim OWF* was omitted and for the variable *s*(*dist OWF*), the number of knots was set to 10. Notably, the meso-scale effects were estimated more reliably in Ref. [31], because the analysis was based on a proper BACI-design intended to measure these effects, which was not the aim of the present study.

**Figure S4.**
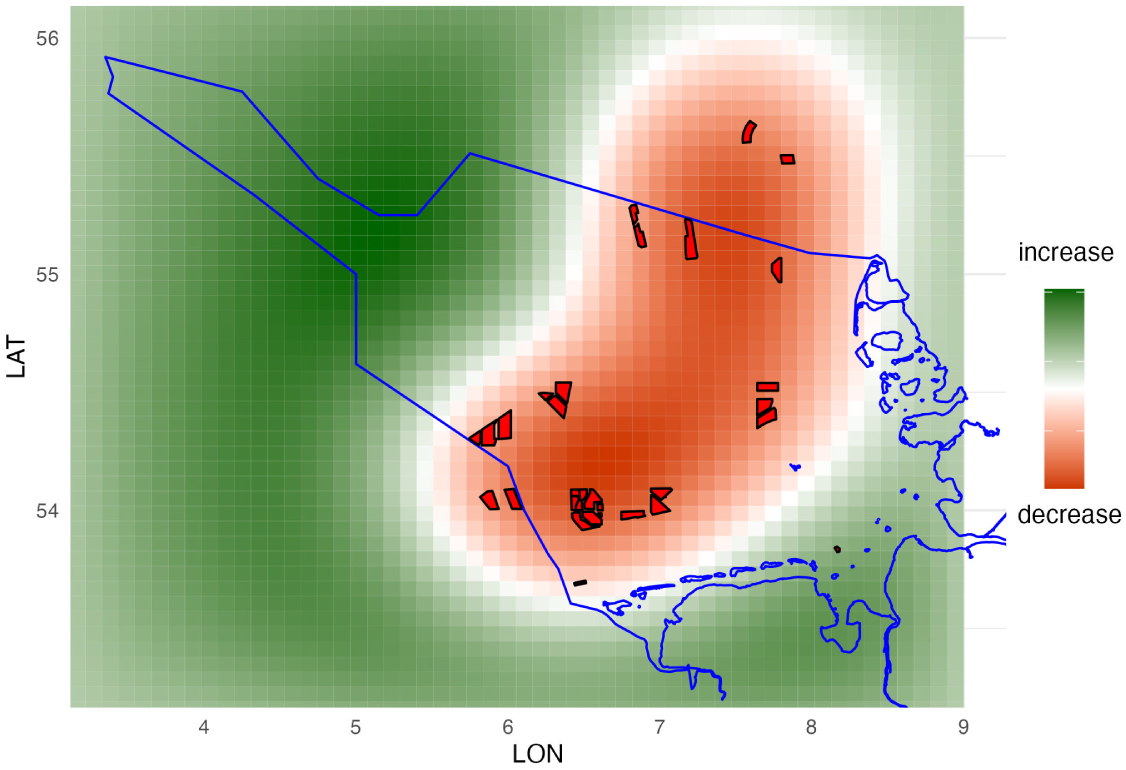
Finally selected PDE-based (large-scale) redistribution patterns (regression variable *sim OWF*) due to current OWF activity (i.e., without any expansions) in the German EEZ for autumn.

## References

[1] Jesse S. Lewis, Susan Spaulding, Heather Swanson, William Keeley, Ashley R. Gramza, Sue Vande- Woude, and Kevin R. Crooks. Human activity influences wildlife populations and activity patterns: Implications for spatial and temporal refuges. Ecosphere, 12(5):e03487, 2021.

[2] Joseph Langridge, Romain Sordello, and Yorick Reyjol. Outcomes of wildlife translocations in protected areas: What is the type and extent of existing evidence? A systematic map protocol. Environmental Evidence, 9(1):16, July 2020.

[3] Moritz Mercker and Klaus Jödicke. Beyond BACI: Offsetting carcass numbers with flight intensity to improve risk assessments of bird collisions with power lines. Ecology and Evolution, 11(23):16716– 16726, 2021.

[4] Neil H. Carter, Simon A. Levin, and Volker Grimm. Effects of human-induced prey depletion on large carnivores in protected areas: Lessons from modeling tiger populations in stylized spatial scenarios. Ecology and Evolution, 9(19):11298–11313, 2019.

[5] Rocío Tarjuelo, Isabel Barja, Manuel B. Morales, Juan Traba, Ana Benítez-López, Fabián Casas, Beatriz Arroyo, M. Paula Delgado, and Francois Mougeot. Effects of human activity on physiological and behavioral responses of an endangered steppe bird. Behavioral Ecology, 26(3):828–838, May 2015.

[6] Susannah S. French, Manuela González-Suárez, Julie K. Young, Susan Durham, and Leah R. Gerber. Human Disturbance Influences Reproductive Success and Growth Rate in California Sea Lions (Zalophus californianus). PLoS ONE, 6(3):e17686, March 2011.

[7] S. Sveegaard, J. Nabe-Nielsen, K. J. Stæhr, T. F. Jensen, K. N. Mouritsen, and J. Teilmann. Spatial interactions between marine predators and their prey: Herring abundance as a driver for the distributions of mackerel and harbour porpoise. Marine Ecology Progress Series, 468:245–253, 2012.

[8] J. Elith and J. R. Leathwick. Species Distribution Models: Ecological Explanation and Prediction Across Space and Time. Annual Review of Ecology, Evolution, and Systematics, 40:677–697, 2009.

[9] Michael Kearney and Warren Porter. Mechanistic niche modelling: Combining physiological and spatial data to predict species’ ranges. Ecology Letters, 12(4):334–350, 2009.

[10] Tyler G. Evans, Sarah E. Diamond, and Morgan W. Kelly. Mechanistic species distribution modelling as a link between physiology and conservation. Conservation Physiology, 3(1):cov056, January 2015.

[11] Philipp Schwemmer and Stefan Garthe. At-sea distribution and behaviour of a surface-feeding seabird, the lesser black-backed gull Larus fuscus, and its association with different prey. Marine Ecology Progress Series, 285:245–258, January 2005.

[12] Brian D. Haig. From Nuisance Variables to Explanatory Theories: A Reformulation of the Third Variable Problem. Educational Philosophy and Theory, 24(2):78–97, January 1992.

[13] E. E. Holmes, M. A. Lewis, J. E. Banks, and R. R. Veit. Partial Differential Equations in Ecology: Spatial Interactions and Population Dynamics. Ecology, 75(1):17–29, 1994.

14. Sergei Petrovski. Partial Differential Equations in Ecology: 80 Years and Counting. https://www.abebooks.com/Partial-Differential-Equations-Ecology-Years-Counting/30885481263/bd, 2021.

15. Volker Grimm and Steven Railsback, F. Individual-Based Modeling and Ecology - Princeton Series in Theoretical and Computational Biology. Princeton University Press, 2005.

[16] Richard A. Stillman. MORPH—An individual-based model to predict the effect of environmental change on foraging animal populations. Ecological Modelling, 216(3):265–276, September 2008.

[17] Bettina Mendel, Philipp Schwemmer, Verena Peschko, Sabine Müller, Henriette Schwemmer, Moritz Mercker, and Stefan Garthe. Operational offshore wind farms and associated ship traffic cause profound changes in distribution patterns of Loons (Gavia spp.). Journal of environmental management, 231:429–438, 2019.

[18] Raul Vilela, Claudia Burger, Ansgar Diederichs, Georg Nehls, Fabian Bachl, Lesley Szostek, Anika Freund, Alexander Braasch, Jochen Bellebaum, Brian Beckers, and Werner Piper. Divers (Gavia spp.) in the German North Sea:Changes in Abundance and Effects of Offshore Wind Farms A study into diver abundance and distributionbased on aerial survey data in the German North Sea. Abschlussbericht Prepared for Bundesverband der Windparkbetreiber Offshore e.V., 2020.

[19] Verena Peschko, Bettina Mendel, Sabine Müller, Nele Markones, Moritz Mercker, and Stefan Garthe. Effects of offshore windfarms on seabird abundance: Strong effects in spring and in the breeding season. Marine Environmental Research, 162:1–12, 2020.

[20] M. Mercker, V. Dierschke, K. Camphuysen, A. Kreutle, N. MArkones, N. Vanermen, and S. Garthe. An indicator for assessing the status of marine-bird habitats affected by multiple human activities: A novel statistical approach. Ecological Indicators, 130:1–14, 2021.

[21] Verena Peschko, Bettina Mendel, Moritz Mercker, Jochen Dierschke, and Stefan Garthe. Northern gannets (Morus bassanus) are strongly affected by operating offshore wind farms during the breeding season. Journal of environmental management, 279:1–11, 2021.

[22] S. Garthe, H. Schwemmer, S. Müller, V. Peschko, N. Markones, and M. Mercker. Seetaucher in der Deutschen Bucht: Verbreitung, Bestnde und Effekte von Windparks, 2018.

[23] M. Mercker. Trend- and population estimates of Gaviidae: Methodological overview. Technical report - published online: https://www.ftz.uni-kiel.de, pages 1–14, 2019.

[24] Michael R. Kearney, Brendan A. Wintle, and Warren P. Porter. Correlative and mechanistic models of species distribution provide congruent forecasts under climate change. Conservation Letters, 3(3):203–213, 2010.

[25] Carsten F. Dormann, Stanislaus J. Schymanski, Juliano Cabral, Isabelle Chuine, Catherine Graham, Florian Hartig, Michael Kearney, Xavier Morin, Christine Römermann, Boris Schröder, and Alexander Singer. Correlation and process in species distribution models: Bridging a dichotomy. Journal of Biogeography, 39(12):2119–2131, 2012.

[26] A. Townsend Peterson, Monica Papeş, and Jorge Soberón. Mechanistic and Correlative Models of Ecological Niches. European Journal of Ecology, 1(2):28–38, December 2015.

[27] Thibaud Rougier, Géraldine Lassalle, Hilaire Drouineau, Nicolas Dumoulin, Thierry Faure, Guillaume Deffuant, Eric Rochard, and Patrick Lambert. The Combined Use of Correlative and Mechanistic Species Distribution Models Benefits Low Conservation Status Species. PLOS ONE, 10(10):e0139194, October 2015.

[28] BSH (2021). Anlage zur Verordnung ueber die Raumordnung in der deutschen ausschließlichen Wirtschaftszone in der Nordsee und in der Ostsee vom 19. August 2021. Anlagenband zum Bundesgesetzblatt Teil I Nr. 58 vom 26. August 2021.

[29] Moritz Mercker, Nele Markones, Kai Borkenhagen, Henriette Schwemmer, Johannes Wahl, and Stefan Garthe. An Integrated Framework to Estimate Seabird Population Numbers and Trends. The Journal of Wildlife Management, 85(4):751–771, May 2021.

[30] S. Garthe, N. Sonntag, P. Schwemmer, and V. Dierschke. Estimation of seabird numbers in the German North Sea throughout the annual cycle and their biogeographic importance. Vogelwelt, 128:163–178, 2007.

[31] Verena Peschko, Henriette Schwemmer, Moritz Mercker, Nele Markones, Kai Borkenhagen, and Stefan Garthe. Cumulative effects of offshore wind farms on common guillemots (Uria aalge) in the southern North Sea - climate versus biodiversity? Biodiversity and Conservation, 33(3):949–970, March 2024.

[32] David C. Schneider. Seabirds and fronts: A brief overview. Polar Research, 8(1):17–21, 1990.

[33] Stefan Garthe, Henriette Schwemmer, Verena Peschko, Nele Markones, Sabine Müller, Philipp Schwemmer, and Moritz Mercker. Large-scale effects of offshore wind farms on seabirds of high conservation concern. Scientific Reports, 13(1):4779, April 2023.

[34] Evelyn F. Keller and Lee A. Segel. Model for chemotaxis. Journal of Theoretical Biology, 30(2):225– 234, February 1971.

[35] Moritz Mercker, Philipp Schwemmer, Verena Peschko, Leonie Enners, and Stefan Garthe. Analysis of local habitat selection and large-scale attraction/avoidance based on animal tracking data: Is there a single best method? Movement Ecology, 9(1):20, 2021.

[36] Arthur N. Popper, Lyndie Hice-Dunton, Edward Jenkins, Dennis M. Higgs, Justin Krebs, Aran Mooney, Aaron Rice, Louise Roberts, Frank Thomsen, Kathy Vigness-Raposa, David Zeddies, and Kathryn A. Williams. Offshore wind energy development: Research priorities for sound and vibration effects on fishes and aquatic invertebrates. The Journal of the Acoustical Society of America, 151(1):205–215, January 2022.

[37] T. Hastie and R. J. Tibshirani. Generalized Additive Models. London, UK: Chapman and Hall, 1990.

[38] S. Wood. Generalized Additive Models: An Introduction with R. Chapman & Hall/CRC, 2017.

[39] A. F. Zuur, E. N. Ieno, N. J. Walker, A. A. Saveliev, and G. M. Smith. *Mixed Effect Models and Extensions in Ecology with R*. Springer Science+Business Media, LLC, New York, 2009.

[40] F. Korner-Nievergelt, T. Roth, S. von Felten, J. Guelat, B. Almasi, and P. Korner-Nievergelt. Bayesian Data Analysis in Ecology Using Linear Models with R, BUGS, and Stan. Elsevier, London, 2015.

[41] H. Akaike. Information theory and an extension of the maximum likelihood principle. *International Sympossium on Information Theroy*, Second Edition:267–281, 1973.

[42] Linden Andreas and Maentyniemi Samu. Using the negative binomial distribution to model overdispersion in ecological count data. Ecology, 92(7):1414–1421, 2011.

[43] C. C. Kokonendji, C. G. B. Demetrio, and S. Dossou-Gbete. Overdispersion and Poisson-Tweedie exponential dispersion models. Monographie del Seminaro Matematico Garcia de Galdeano, 31:365– 374, 2004.

[44] A. F. Zuur and E. N. Ieno. A protocol for conducting and presenting results of regression-type analyses. Methods Ecol Evol 7: 636645, 2016.

45. R. Core Team. R: A language and environment for statistical computing. R Foundation for Statistical Computing, Vienna, Austria. URL https://www.R-project.org. 2022.

46. Soetaert Karline, Setzer R. Woodrow, and Petzoldt Thomas. Solving Differential Equations in R. The R Journal, 1281(1):31–34, September 2010.

[47] Pebesma, E.J., R. S. Bivand. Classes and Methods for Spatial Data: The sp Package. R News, pages 1–21, 2005.

[48] E. J. Pebesma. Multivariable geostatistics in S: The gstat package. Computers & Geosciences, 30(7):683–691, 2004.

49. Roger Bivand. Rgdal: Bindings for the ‘Geospatial’ Data Abstraction Library version 1.5-32 from CRAN. https://rdrr.io/cran/rgdal/.

[50] Robert J. Hijmans, Charles Karney (GeographicLib), Ed Williams, and Chris Vennes. Geosphere: Spherical Trigonometry, October 2021.

[51] Joseph Larmarange. prevR: Estimating Regional Trends of a Prevalence from a DHS and Similar Surveys, May 2022.

[52] Roger Bivand, Colin Rundel, Edzer Pebesma, Rainer Stuetz, and Karl Ove Hufthammer. Rgeos: Interface to Geometry Engine - Open Source (‘GEOS’), October 2017.

[53] H. Wickham. Ggplot2: Elegant Graphics for Data Analysis. Springer-Verlag New York, 2009.

[54] Verena Peschko, Moritz Mercker, and Stefan Garthe. Telemetry reveals strong effects of offshore wind farms on behaviour and habitat use of common guillemots (Uria aalge) during the breeding season. Marine Biology, 167(118):1–13, 2020.

[55] Todd W. Arnold. Uninformative Parameters and Model Selection Using Akaike’s Information Criterion. The Journal of Wildlife Management, 74(6):1175–1178, 2010.

